# Reln haploinsufficiency enhances fentanyl-induced locomotion and striatal activity without affecting opioid reinforcement and relapse-like behavior

**DOI:** 10.64898/2026.02.21.707172

**Authors:** Carl Litif, Avraham M. Libster, Shane Desfor, Tanieng Huang, Leanne Liaw, Annika Cheng, Francesca Telese

**Affiliations:** Department of Psychiatry, School of Medicine, University of California, San Diego

## Abstract

The *Reln* gene encodes the extracellular glycoprotein Reelin that regulates synaptic plasticity and activity-dependent gene expression with implications in several neuropsychiatric disorders, including substance use disorder. While reduced *Reln* expression alters responses to psychostimulants and cannabinoid, its role in opioid-related behaviors remains unknown. Here, we examined whether *Reln* haploinsufficiency modifies behavioral and molecular responses to the synthetic opioid fentanyl. Heterozygous Reeler (*Reln*+/−) mice and wild-type littermates were assessed using using complementary contingent and non-contingent models of fentanyl exposure, including multi-phase fentanyl intravenous self-administration paradigm, conditioned place preference paradigm, locomotor assay, and dorsal striatal immediate early gene expression. *Reln* haploinsufficiency did not alter acquisition, extinction, or cue-induced reinstatement during self-administration, indicating stable opioid reinforcement and relapse-like behavior. Progressive ratio testing revealed a sex-dependent effect in which male *Reln*+/− mice showed reduced motivation for fentanyl compared to male wild-type mice. In contrast, following passive fentanyl exposure, *Reln*+/− mice exhibited enhanced fentanyl-induced locomotion and increased Fos immunoreactivity in the dorsal striatum, while CPP remained unchanged. Together, these findings demonstrate that *Reln* haploinsufficiency does not substantially modify opioid reinforcement or cue-driven drug seeking but enhances acute pharmacological sensitivity to fentanyl. These results identify *Reln* as a modulatory factor in opioid-responsive neural circuits that preferentially influences acute drug-evoked neuronal activation rather than the associative learning processes underlying opioid reinforcement.

## Introduction

Opioid use disorder (OUD) affects millions globally (Degenhardt et al., 2019; Hser et al., 2001) and is characterized by chronic drug intake, relapse vulnerability, and maladaptive learning processes involving the encoding of drug-associated cues and contexts (Everitt & Robbins, 2005). These behaviors reflect molecular and cellular perturbations within corticostriatal circuits, particularly the dorsal striatum, which supports action-selection, habit-formation, and cost-benefit valuation of reward-seeking behaviors (Cucinello-Ragland et al., 2026). Opioids, including the highly potent synthetic agonist fentanyl, robustly engage striatal signaling pathways that regulate synaptic plasticity, neuronal excitability, and immediate early gene (IEG) expression (Bisagno & Cadet, 2019). Although opioid-induced behavioral adaptations have been extensively characterized, the molecular regulators that govern these plasticity processes remain poorly defined.

Reelin (*Reln*) is a large extracellular glycoprotein (D’Arcangelo et al., 1997) expressed mainly in the striatum, cortex, and cerebellum (Ikeda & Terashima, 1997). While best known for its developmental role in neuronal migration, Reelin is increasingly recognized for its functions in adult neuronal plasticity through modulation of IEG expression (G. H. Lee et al., 2014; Telese et al., 2015), dendritic spines regulation (Bosch et al., 2016; H.-J. Lee et al., 2023; Niu et al., 2008; Rogers et al., 2011), and long-term potentiation (Beffert et al., 2005; Qiu et al., 2006) in the adult brain. In humans, altered RELN expression has been implicated in several neuropsychiatric disorders characterized by disrupted synaptic plasticity (Fatemi et al., 2000), including autism (Wang et al., 2014; Zhang et al., 2002), schizophrenia (Costa et al., 2001; Grayson et al., 2005), Alzheimer’s disease (Cuchillo-Ibañez et al., 2016; Seripa et al., 2008), and attention-deficit/hyperactivity disorder (Chen et al., 2017).

Emerging preclinical evidence also links Reelin signaling to phenotypes associated with substances of abuse. Experimental reduction of *Reln* expression in mice modifies behavioral and cellular responses to cannabinoids (Iemolo et al., 2021; Silva-Hurtado et al., 2024; Zuo et al., 2025) and psychostimulants (Brida et al., 2025; de Guglielmo et al., 2022; Matsuzaki et al., 2007), including drug-induced locomotion, reward-related learning, and striatal activation patterns. In parallel, clinical studies report associations between altered RELN expression and phenotypes associated with alcohol use disorder (Escudero et al., 2023; Legaki et al., 2024). Together, these findings suggest that Reelin signaling may influence how drug experiences are encoded within reward-related neural circuits.

Despite these advances, the role of Reelin in opioid-related behaviors remains unknown. Opioids differ from psychostimulants and cannabinoids in both receptor mechanisms, yet share aspects of circuit recruitment for striatal neuronal pathways involved in reinforcement learning and adaptive responding to changing reward contingencies (Le Merrer et al., 2009). Given the role of Reelin signaling in regulating activity-dependent transcription and synaptic plasticity, alterations in Reelin signaling may shape how opioid exposure engages striatal neuronal activation and reinforcement-related learning. To address this gap, we evaluated heterozygous Reeler (*Reln*+/−) mice and wild-type (WT) littermates (D’Arcangelo et al., 1995) in complementary contingent and non-contingent models of fentanyl exposure. Using a multi-phase fentanyl intravenous self-administration (IVSA) paradigm, we examined acquisition, motivation, extinction, and cue-induced reinstatement to determine whether *Reln* haploinsufficiency alters opioid reinforcement or relapse-like behavior. In parallel, we examined conditioned place preference (CPP), fentanyl-induced locomotion, and Fos immunoreactivity in the dorsal striatum to assess opioid-evoked neuronal activation under passive exposure to fentanyl. Altogether, this study dissociates the role of Reelin in reinforcement processes from adaptive responding associated with fentanyl exposure.

## Materials and Methods

### Mice

All animal care and experimental procedures were approved by the UCSD Institutional Animal Care and Use Committee. Male and female *Reln*+/− and WT littermate controls were bred in house using the B6C3Fe a/a-Relnrl/J line (The Jackson Laboratory, #000235). Genotypes were determined by PCR-based analysis of tail biopsies (de Guglielmo et al., 2022). Mice were group-housed (2-4 per cage), except during IVSA where mice were individualized immediately following catheterization, under a 12-h light/dark cycle with ad libitum food and water except during behavioral testing where mice were maintained on a reverse light cycle. Mice were handled for a minimum of three days before the start of each experiment. For multi-phase IVSA, n = 8 WT and n = 5 *Reln*+/− male mice, and n= 5 WT and n = 8 *Reln*+/− female mice were used across two cohorts. For CPP and locomotion, n = 4 WT and n = 4 *Reln*+/− male, and n = 4 WT and n = 4 Reln+/− female mice were used per low or high dose. For Fos immunohistochemistry n = 4 WT and n = 4 *Reln*+/− male mice were used for the low dose. Across all experiments, mice ranged from 12 to 16 weeks of age at the beginning of testing.

### Drug Preparation

Powdered fentanyl citrate (Cayman Chemical Company) was dissolved in 0.9% saline to a stock concentration of 5 mg/ml. For IVSA, 5 mg/ml stock of fentanyl citrate and saline was further diluted to a working solution with saline for a final concentration of 0.025 mg/ml to allow a dynamic range of 15-1.5 ug/kg/infusion that was dependent on the duration of infusion. For acute exposure and CPP, 5 mg/ml stock of fentanyl citrate and saline was further diluted to a working solution with saline for a final concentration of 0.003 or 0.012 mg/ml to allow for 20 µg/kg and 80 µg/kg IP injections, respectively.

### Fentanyl IVSA

For IVSA procedures, mice were individually housed to prevent cagemates from damaging the back-mounted catheters. Mice were food-restricted to 80% of their normal intake to increase motivation for operant conditioning (Chevée et al., 2023).

#### Surgical procedures

Mice underwent intravenous jugular catheterization using standard procedure (Margiani et al., 2022; Valles et al., 2022). Following surgery, mice were given daily post-operative care with intravenous flushes of heparinized saline containing cefazolin antibiotic to prevent clogging and infection. Once mice recovered to pre-surgical body weight and had a dry wound area (3-5 days), the mice underwent fentanyl IVSA.

#### Acquisition (FR1, 15 µg/kg/infusion)

Self-administration was conducted in standard operant-conditioning chambers (Med Associates) equipped with two retractable levers positioned on one wall where an active lever was reinforced with intravenous delivery of fentanyl as opposed to an inactive lever that featured no consequences. At the start of each IVSA session, the white house lights were illuminated at the top of the arena, mice were given a passive infusion of fentanyl at the dose they would receive for the session, and a yellow availability light cue above the active lever was illuminated. Upon interaction with the active lever, the yellow availability light cue extinguished illumination, fentanyl was delivered, and a pure tone was played for 10 seconds. Active lever presses within this 10 second tone did not allow another infusion delivery. Following the 10 second tone, the availability light was illuminated again to signal another fentanyl infusion is available. All active or inactive lever presses and the infusions were recorded. The house lights were illuminated with both active and inactive levers extended for the entire session. Each session was 2 hours in duration and conducted daily.

#### De-escalating Dose-Response Testing

Following acquisition, mice underwent dose-response testing in which fentanyl unit dose was sequentially reduced (15, 10, 5, 1.5 µg/kg/infusion), with two consecutive 2-hour sessions at each dose under an FR1 schedule. Dose order was identical for all animals. Different IVSA doses were implemented by adjusting the duration of infusions (0.1 mg/sec) in the MedPC software while using the same concentration of fentanyl (0.25 mg/ml; Supplemental Table 1). Dose-response sessions followed the same cue presentation pattern. Mice were given an initial passive infusion of fentanyl at the start of the session with the dose they would receive for the session. Infusion number, active and inactive lever presses, and total intake were recorded.

#### Dose Reset Testing

Following dose-response testing, two “dose reset” sessions were conducted at 15 µg/kg/infusion dose to re-stabilize responding to the acquisition dose.

#### Progressive Ratio Testing

Following dose-response and dose-reset sessions, motivation was assessed under a linear PR schedule conducted at the lowest unit dose (1.5 µg/kg/infusion). Under the PR schedule, the response requirement increased incrementally on a linear scale for each subsequent infusion (1, 2, 3, 4, …). The breakpoint was defined as the highest completed ratio resulting in infusion delivery. Active and inactive lever presses, infusions, and breakpoints were recorded.

#### Extinction and Cue-Induced Reinstatement

Following PR testing, mice underwent extinction training in which lever presses had no programmed consequences (no fentanyl delivery or cues). Extinction sessions were 2 hours and conducted daily for 10 sessions. A 30-minute cue-induced reinstatement session was conducted by reintroducing the previously fentanyl-paired cues without drug delivery. Lever presses during reinstatement were compared to the first 30 minutes of the final extinction session.

### RT-qPCR

Following cue-induced reinstatement, mice were euthanized immediately after the session termination. Dorsal striatal tissue punches (+1.5-0.0 mm bregma) were collected and sectioned using a vibratome (Leica), flash frozen and stored at −80C. Total RNA was extracted from frozen tissues using Direct-Zol RNA Miniprep (Zymo Research, #R2050). cDNA was synthesized from 100 ng RNA using the Maxima H Minus First Strand cDNA Synthesis Kit with dsDNase (Thermo Fisher Scientific, #K1682). RT-qPCR was performed using the following primer sequences (Supplemental Table 1) and SsoAdvanced Universal SYBR Green Supermix (Bio-Rad, #1725270). Gene expression was normalized to the housekeeping gene, *Gapdh*, using the 2^-ΔCt method (Livak & Schmittgen, 2001).

### Fentanyl CPP

CPP was conducted in a two-compartment apparatus with distinct visual and tactile cues. On the pre-test day, mice were allowed free access to both chambers for 30 minutes to assess baseline contextual preference. A counter-biased design was used. If a mouse displayed over 55% preference for one chamber during the pre-test session, the least preferred chamber was assigned as the fentanyl paired context. If no strong baseline preference was observed (≤55%), pairing assignment was counterbalanced across genotypes. Fentanyl conditioning consisted of eight total treatment days where treatments alternated daily (four fentanyl, four saline in opposite chambers). Mice were treated with fentanyl in two separate cohorts, a low dose (20µg/kg) and a high dose (80µg/kg). On saline-pairing days, mice received an equivalent volume of saline as they would on fentanyl-treated days and were confined to the alternate chamber. On the test day, mice were allowed free access to both compartments for 30 minutes in a drug-free state.

All sessions were 30 minutes in length and recorded using red lighting and a camera positioned above the apparatus. CPP videos were analyzed using DeepLabCut v3.0 to track estimated position of the mouse midpoint across frames using an 8-pointed pose-estimation framework. The mouse was scored as occupying a chamber when its centroid fell within that chamber’s boundary. Time spent in each chamber was calculated as the number of frames assigned to each chamber divided by the video frame rate (30 frames per second). Videos were visually inspected to confirm accurate chamber assignments. Preference score was calculated as the change in percentage of time spent in the fentanyl-paired chamber compared to the saline-paired chamber between the pre-test and test sessions.

### Locomotion Analysis

Locomotor activity during fentanyl exposure was quantified using DeepLabCut v3 to track the mouse location across frames using 8-pointed pose-estimation. The total distance (meter) traveled was calculated as the total sum of the difference between the animal’s centroid position in consecutive frames. The reported average speed (meter / minute) was calculated by dividing the total distance by the total time in the session (30 minutes). All analyses were performed using custom Python and Matlab (Mathworks) scripts and were conducted blind to genotype.

### Acute Fentanyl Exposure and Fos Immunostaining

In a separate cohort, mice received an acute fentanyl injection (20 µg/kg, i.p.) and were perfused in 1x PBS followed by 4% PFA 90 minutes later. Brains were post-fixed (overnight in 4°C, 4% PFA), cryoprotected (30% sucrose solution, until sinking), and sectioned coronally using a microtome (Microm HM440E). Three mounted sections (20um) featuring the dorsal striatum (bregma 0.74 mm) per each genotype were used. Immunohistochemistry (IHC) for Fos protein was performed as follows: 1-hour incubation in blocking buffer [1x PBS (Millipore Sigma; # P4244-100ML), 10% normal goat serum (Millipore Sigma; # S26-M), 0.3% TritonX-100 (Millipore Sigma; #X100-5ML)] at room temperature followed by overnight incubation in Fos primary antibody dilution solution [1x PBS, 5% normal goat serum, 0.1% TritonX-100, and 1:2500 anti-Fos rabbit (Synaptic Systems; # 226 008)] at 4°C, three 15-minute washes in washing buffer (1x PBS, 0.3% TritonX-100), 2-hour incubation in secondary antibody solution [1x PBS, 5% normal goat serum, 0.1% TritonX-100, and 1:500 Donkey anti-Rabbit AlexaFluor-680 (ThermoFisher; # A10043)] at room temperature, three 15-minute washes in washing buffer at room temperature, 30-minute incubation in 1x PBS with 600 nM DAPI at room temperature, two 15-minute washes in washing buffer at room temperature. Sections were coverslipped with mounting medium (Millipore Sigma, # 10981). Images of the dorsal striatum were acquired using a fluorescence microscope (Keyence; BZ-X810) under identical exposure settings across animals. Cropped region-of-interest (ROI) images corresponding to the dorsal striatum were generated in ImageJ (Schneider et al., 2012) using standardized anatomical boundaries of the corpus collosum, lateral ventricle, and anterior commissure. Automated cell quantification was performed using CellProfiler (Carpenter et al., 2006) to identify DAPI-positive nuclei and Fos-positive nuclei. For each mouse, three coronal slices (six total sections including left and right hemispheres) were analyzed. The average number of Fos+/DAPI+ cells per section was calculated and used as the technical replicate for statistical analysis.

### Statistical Analysis

All statistical analyses and data visualization were conducted in R (version 4.5.0). For IVSA, separate models were fitted for each experimental phase (acquisition, dose-response, dose reset, progressive ratio, extinction, and cue-induced reinstatement). IVSA count data (lever presses or infusions) were analyzed using negative-binomially-distributed generalized linear mixed-effects models (GLMMs) after failing to meet Gaussian assumptions while featuring overdispersion. IVSA continuous, positive data (total intake) were analyzed using gamma-distributed GLMMs after failing to meet Gaussian assumptions while featuring overdispersion. For lever presses, infusions, or total intake responses, fixed effects included genotype, lever, day, session condition, sex, with mice included as a random effect. For breakpoint response, fixed effects included genotype and sex. For GLMMS, fixed effects and interactions were evaluated and reported using likelihood ratio tests (LRTs). Post-hoc pairwise comparisons were performed using estimated means with false discovery rate (FDR) correction. RT-qPCR relative expression values were modeled with linear mixed models (LMM) after meeting Gaussian assumptions to compare the fixed effects of genotype and sex for each gene of interest, where mice and their nested qPCR technical replicates were considered random effects. CPP data, which met Gaussian assumptions, were analyzed using a LMM to assess fixed effects of genotype, dose, and change in preference for the fentanyl-paired chamber, using sex as a covariate effect, and mice as a random effect. Similarly, fentanyl-induced locomotion (average speed) was analyzed using an LMM including genotype and dose as fixed effects, sex as a covariate effect. For CPP and locomotion LMMs, models that included sex as interaction term did not find any significant interactions with sex; given the small sample size we used simplified LMMs that included sex as covariate. The percentage of Fos-positive nuclei relative to total DAPI-positive nuclei was modeled with a LMM to compare the fixed effect of genotype, where mice and their nested technical replicated slices were considered random effects. For LMMs, fixed effects and interactions were evaluated and reported using analysis of variance (ANOVA). Statistical significance for LRTs and ANOVAs were defined as p < 0.05 while a FDR adjusted adjP < 0.05 was used for post hoc pairwise comparisons. Exact test statistics and degrees of freedom are reported in Supplemental Table 2-6.

## Results

### *Reln* haploinsufficiency does not alter the acquisition of fentanyl

To determine whether reduced *Reln* expression alters opioid reinforcement or relapse-like responding, *Reln*+/− and wild-type (WT) littermates were assessed in a multi-phase fentanyl IVSA paradigm, including acquisition, dose-response, dose reset, progressive ratio (PR), extinction, and cue-induced reinstatement (Figure 1A).

**Figure 1.**
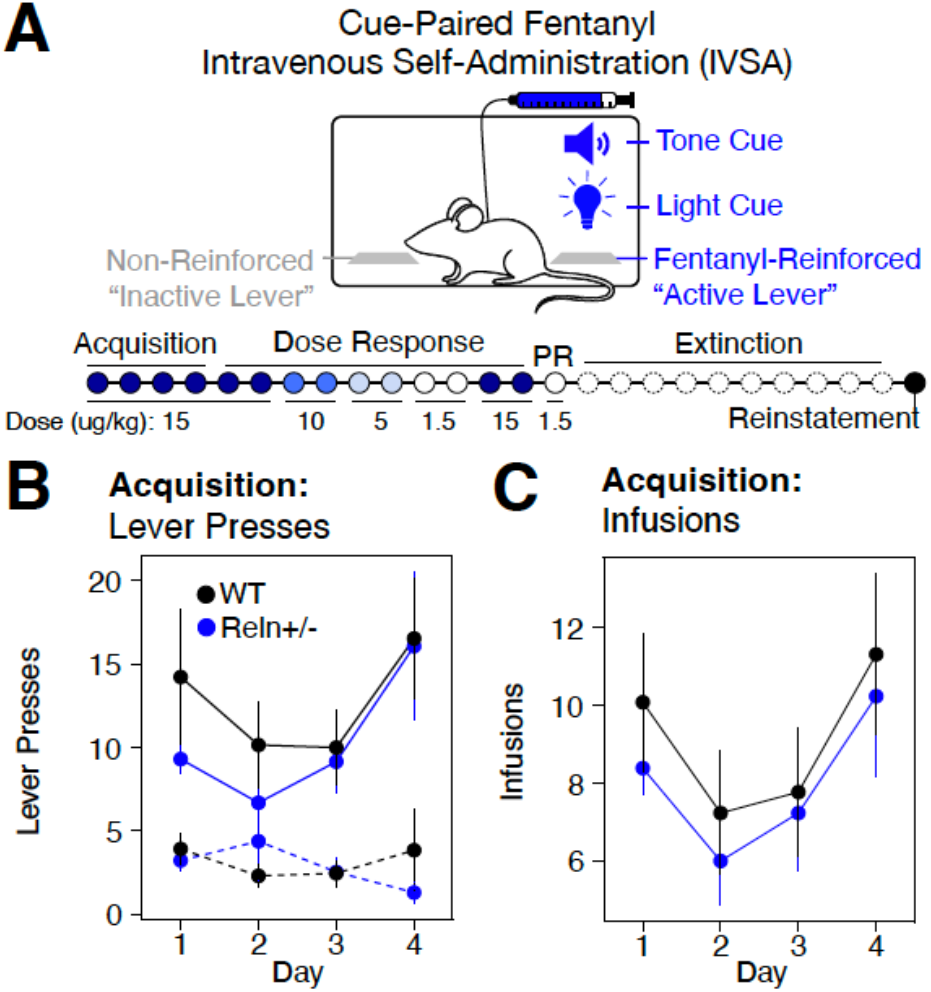
Assessing *Reln* haploinsufficiency within a fentanyl IVSA paradigm. **(A)** Schematic of fentanyl IVSA comparing *Reln*+/−(n=13) and WT (n=13) mice. **B-C** Active (solid line) and inactive (dashed line) lever presses (**B**) or fentanyl infusions (**C**) across acquisition sessions (Days 1-4), collapsed across sex. Data are mean ± SEM.

During a four-day acquisition phase (15 ug/kg/infusion), both genotypes acquired fentanyl self-administration, as evidenced by a significant main effect of lever (χ^2^(1) = 10.823, p = 0.001, Supplemental Table 2.01), indicating that mice across both genotypes preferentially pressed the active over the inactive lever. This lever discrimination persisted across sessions, reflected in a significant lever × day interaction (χ^2^(3) = 9.911, p = 0.019, Supplemental Table 2.01). Importantly, there was no main effect of genotype on lever presses or infusions during acquisition (Figure 1B-C, Supplemental Table 2.01-2.02), indicating that baseline fentanyl reinforcement learning was preserved in *Reln*+/− mice.

Together, these data indicate that *Reln* haploinsufficiency does not modify baseline opioid reinforcement.

### Genotype-dependent differences emerge during fentanyl dose manipulation

To determine whether *Reln* haploinsufficiency alters dose-dependent fentanyl responding, we implemented a de-escalating dose response phase in which the unit dose was reduced sequentially across sessions (15, 10, 5, 1.5 µg/kg/infusion). When modeled by dose, both genotypes exhibited the expected inverse relationship between unit dose and active lever presses, with responding increasing as the unit dose decreased (Figure 2A; lever x dose = χ^2^(3) = 14.51, p = 0.002, Supplemental Table 2.03). Post hoc comparisons confirmed differences in responding across doses (Supplemental Table 2.03). A significant lever × genotype × sex interaction was detected for active lever presses when modeled by dose (Supplemental Figure 1A; lever × genotype × sex: χ^2^(1) = 6.222, p = 0.013, Supplemental Table 2.03); however, post hoc pairwise comparisons did not detect significant genotype differences within any sex or lever condition after FDR correction (Supplemental Table 2.03), indicating that this interaction did not reflect a consistent genotype-dependent effect and may reflect limited power within sex-by-genotype subgroups. Modeling sessions by individual day similarly confirmed the dose-dependent pattern of lever responding (lever × day: χ^2^(7) = 20.17, p = 0.005; Supplemental Table 2.04), with no significant genotype effects.

**Figure 2.**
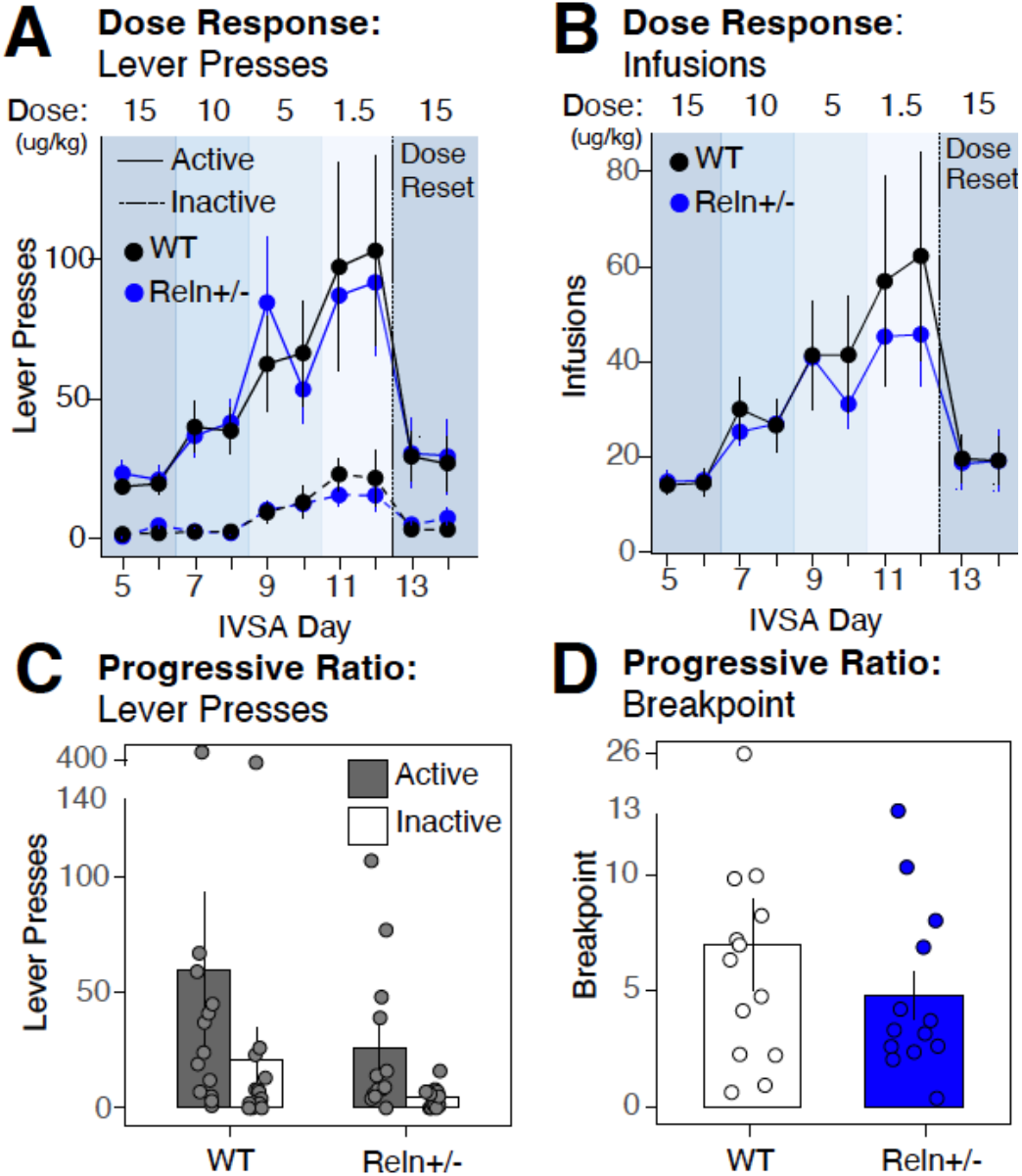
*Reln* haploinsufficiency does not alter fentanyl dose-response or motivation during intravenous self-administration. **A-B** Active (solid line) and inactive (dashed line) lever presses (**A**) or fentanyl infusions (**B**) across the dose-response phase, in which the unit dose was decreased sequentially from 15 to 10, 5, and 1.5 µg/kg/infusion over IVSA Days 5–12, followed by a dose reset to 15 µg/kg/infusion (Day 13-14; dotted vertical line). Data are mean ± SEM. (**C-D**) Total active and inactive lever presses (**C**) and breakpoint (**D**) during the progressive ratio session, in which the response requirement increased with each successive infusion. Data are mean ± SEM. WT, n = 13; Reln+/−, n = 13.

Fentanyl infusions followed a comparable pattern, increasing as unit dose decreased when modeled by dose (Figure 2B; main effect of dose: χ^2^(3) = 36.14, p < 0.001, Supplemental Table 2.05) or by session day (main effect of day: χ^2^(7) = 90.52, p < 0.001, Supplemental Table 2.06), with no genotype effects detected in either model. Total fentanyl intake similarly varied as a function of dose (main effect of dose: χ^2^(3) = 39.62, p < 0.001, Supplemental Table 2.07) and session day (main effect of day: χ^2^(7) = 146.4, p < 0.001, Supplemental Table 2.08), producing an inverted U-shaped function when collapsed across doses (Supplemental Figure 2), again with no genotype differences.

Following dose-response testing, mice were returned to the acquisition dose (15 µg/kg/infusion) for two dose reset sessions. Both genotypes maintained lever discrimination during this phase (Figure 2A; main effect of lever: χ^2^(1) = 17.12, p < 0.001, Supplemental Table 2.09), with no change in responding across the sessions and no genotype differences in lever presses or infusions (Supplemental Tables 2.09–2.10), indicating stable performance at the original acquisition dose.

Together, these findings demonstrate that both genotypes responded appropriately to changes in unit dose and that *Reln* haploinsufficiency did not produce a consistent effect on dose-dependent fentanyl responding.

### *Reln* haploinsufficiency does not alter motivation for fentanyl intake

Following the two dose-reset sessions, motivation for fentanyl was assessed under a linear PR schedule conducted at the lowest dose in the dose-response (1.5 ug/kg/infusion). Mice showed strong lever bias overall (Figure 2C; main effect of lever: χ^2^(1) = 6.964, p = 0.008; Supplemental Table 2.11), with no detectable genotype differences in lever presses (Supplemental Table 2.11). In contrast, during the breakpoint, a significant genotype × sex interaction was detected (Figure 2D; χ^2^(1) = 4.402, p = 0.036; Supplemental Table 2.12). Post hoc comparisons revealed this interaction was driven by two related patterns: male WT mice earned significantly more infusions than male *Reln*+/− mice (Supplemental Figure 1D; estimate = 0.48, z = −2.365, adjP = 0.018; Supplemental Table 2.12), and male WT mice earned significantly more infusions than female WT mice (estimate = 0.48, z = −2.365, adjP = 0.018; Supplemental Table 2.12), whereas no significant difference was detected between female and male *Reln*+/− mice.

Overall, these results suggest that *Reln* haploinsufficiency reduces motivation for fentanyl intake in males to levels comparable to those observed in females of either genotype rather than their male WT counterparts. This pattern points to a sex-dependent role of *Reln* signaling in the effort exerted to obtain opioid reinforcement under conditions of increasing response cost.

### *Reln* haploinsufficiency does not alter extinction or reinstatement of fentanyl self-administration

During extinction, lever discrimination decreased across sessions with no detectable differences between levers or genotype across days (Figure 3A; Supplemental Table 2.13), indicating loss of cue-directed lever pressing without cue presentation or fentanyl delivery for both WT and Reln+/− mice.

**Figure 3.**
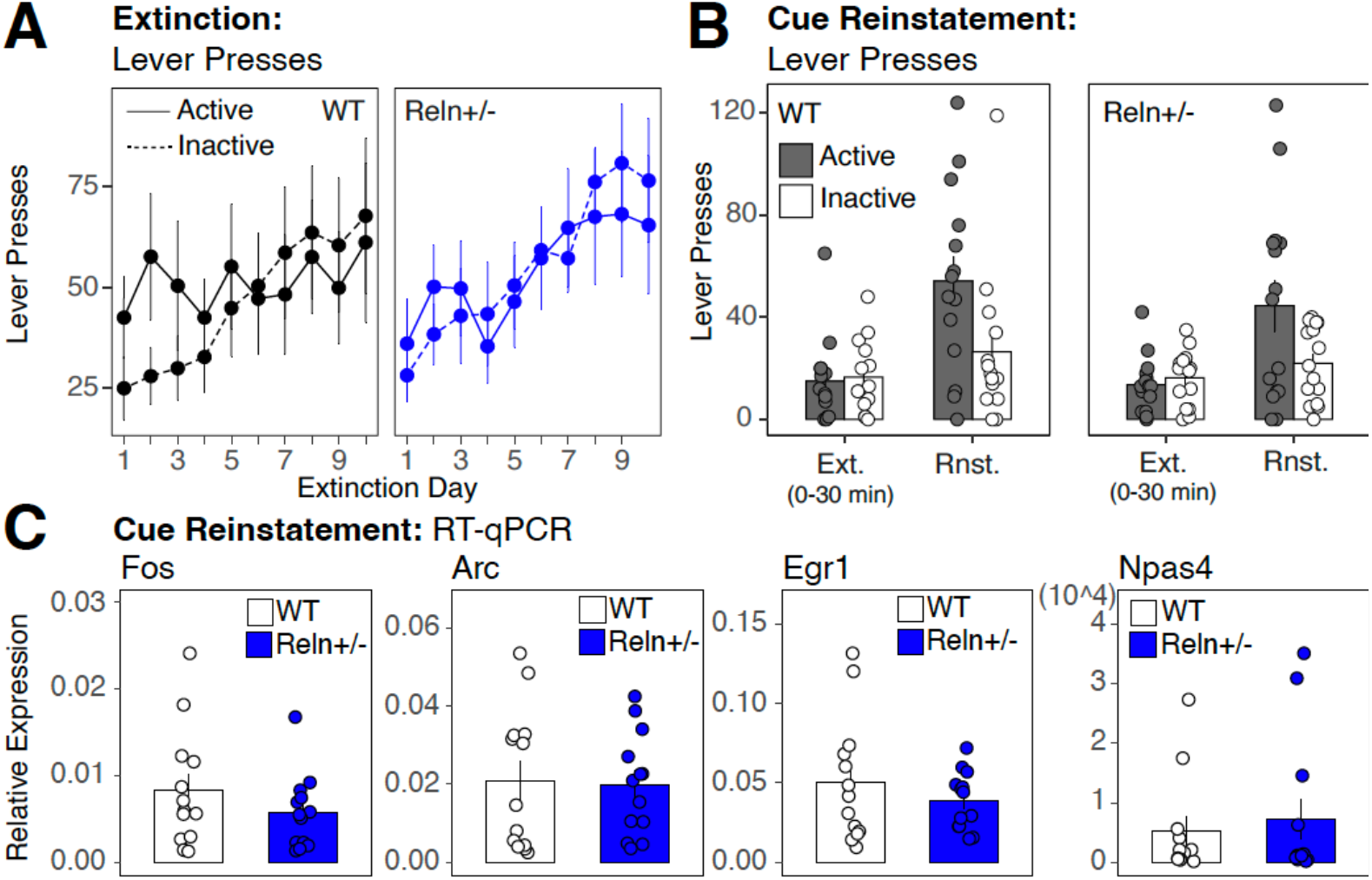
*Reln* haploinsufficiency does not alter extinction, cue-induced reinstatement, and dorsal striatal immediate early gene expression following reinstatement. **(A)** Active (solid line) and inactive (dashed line) lever presses across 10 daily extinction sessions for WT (black) and *Reln*+/− (blue) mice. (**B**) Comparison of active and inactive lever presses during the final 30 minutes of the last extinction session (Ext.) and during the subsequent 30-minute cue-induced reinstatement session (Rnst.). (**C)** Relative expression of immediate early genes *Fos, Arc, Egr1*, and *Npas4* in the dorsal striatum following cue-induced reinstatement, normalized to *Gapdh*, in WT and *Reln*+/− mice. Data are mean ± SEM. WT, n = 13; *Reln*+/−, n = 13.

During cue-induced reinstatement, in the presence of fentanyl-associated cues without fentanyl delivery, both genotypes exhibited robust reinstatement of active lever responding compared to the last extinction session, demonstrated by a significant main effect of session condition (Figure 3B; χ^2^(1) = 14.43, p < 0.0001; Supplemental Table 2.14). A significant session condition × genotype × sex interaction was detected (Supplemental Figure 1; χ^2^(1) = 5.66, p = 0.017; Supplemental Table 2.14), which post hoc analysis indicated was driven by greater active lever presses in WT female compared to WT males specifically during reinstatement (estimate = 2.358, z = 2.023, adjP = 0.043; Supplemental Table 2.14), with no significant genotype differences within either sex for either session condition (Supplemental Table 2.14).

Consistent with the absence of genotypic differences in cue-induced reinstatement behavior, the expression of IEGs (*Fos, Arc, Egr1, Npas4*), time-locked to the reinstatement session in the dorsal striatum did not differ between genotypes (Figure 3C; Supplemental Table 3).

Together, these findings demonstrate that *Reln* haploinsufficiency does not alter cue-induced reinstatement of drug-seeking in either sex. The sex difference observed in reinstatement magnitude among WT mice with females showing greater responding than males was not replicated in *Reln*+/− mice, suggesting a possible interaction between sex and *Reln* expression in the modulation of cue-driven drug-seeking, though the study was not specifically powered to test this.

### *Reln* haploinsufficiency enhances non-contingent acute fentanyl-induced locomotion and dorsal striatal Fos expression without an effect on CPP

To examine opioid-evoked responses independent of instrumental contingencies, we evaluated passive fentanyl exposure using CPP, locomotor analysis, and Fos immunoreactivity in the dorsal striatum as a marker of neuronal activation.

CPP doses (20 or 80 µg/kg, i.p.) were selected to approximate contingent intake observed during IVSA at the lowest and highest unit doses, respectively (Figure 4A). *Reln*+/− mice did not alter fentanyl CPP at either dose (Figure 4B; Supplemental Table 4).

**Figure 4.**
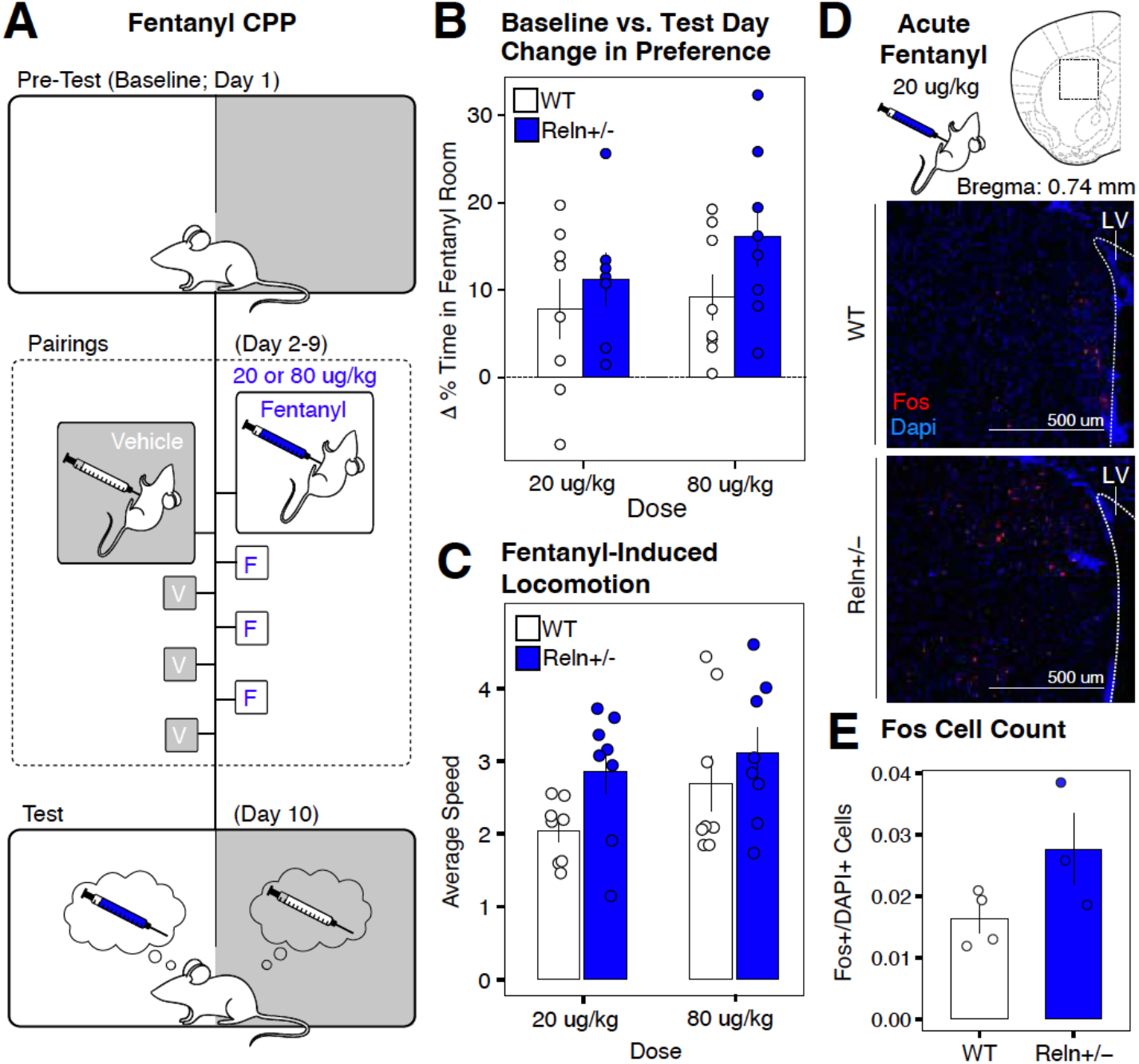
*Reln* haploinsufficiency enhances acute fentanyl-induced locomotion and dorsal striatal Fos immunoreactivity without altering conditioned place preference. **(A)** Schematic of the CPP paradigm. (**B)** Change in percent time spent in the fentanyl-paired room at 20 µg/kg (left) or (80 µg/kg or (right). (**C)** Average locomotor speed (total distance / total time; meter / minute) during the first CPP conditioning session following acute fentanyl injection at 20 µg/kg (left) or 80 µg/kg or (right). (**D)** Representative fluorescent image of the dorsal striatum from WT (top) and *Reln*+/− (bottom) mice following acute injection (20 µg/kg). Fos immunoreactivity is shown in red; DAPI nuclear counterstain is in blue. Scale bar, 500um. (**E)** Quantification of Fos+ cells as a proportion of total DAPI+ cells in the dorsal striatum of a different cohort of mice following acute fentanyl injection (20 µg/kg, i.p.). Data are mean ± SEM. WT, n=4; *Reln*+/−, n=4 mice. *p<0.05.

Locomotor activity was quantified from the first fentanyl conditioning session to quantify the acute pharmacological response to fentanyl before repeated exposure could introduce sensitization or tolerance effects. *Reln+/−* mice showed a greater fentanyl-induced locomotion than WT mice (main effect of genotype: (χ^2^(1) = 5.635, p = 0.018; Supplemental Table 5; Figure 4C). Alongside this genotype difference, we found a dose-dependent difference for fentanyl-induced locomotion (main effect of dose: (χ^2^(1) = 3.995, p = 0.046; Supplemental Table 5), with higher doses producing greater locomotor activation.

However, no significant genotype x dose interaction was detected (Supplemental Table 5), suggesting that *Reln* haploinsufficiency broadly enhances acute behavioral sensitivity to fentanyl independent of the dose. A sex-dependent effect was also detected (main effect of sex: χ^2^(1) = 7.72, p = 0.0056; Supplemental Table 5; Supplemental Figure 3B), with males showing greater locomotion compared to females across genotypes.

To determine whether this behavioral difference was accompanied by altered neuronal activation, a separate cohort of male mice received an acute fentanyl injection at the low dose (20 µg/kg, i.p.), and Fos immunoreactivity was quantified 90 minutes later in the dorsal striatum (Figure 4D). Quantifying the population of Fos-positive (Fos+) cells relative to the total Dapi-positive (Dapi+) cellular population, *Reln*+/− mice showed a significant increase in the fraction of Fos+ cells out of the total cellular population [(Fos+/Dapi+) / Dapi+] compared to WT mice (main effect of genotype: χ^2^(1) = 4.1873, p = 0.041; Figure 4E; Supplemental Table 6).

Together, these findings demonstrate that *Reln* haploinsufficiency enhances acute fentanyl-induced locomotion and neuronal activation in the dorsal striatum, without affecting passively learned reinforcing properties of fentanyl.

## Discussion

This study provides the first characterization of *Reln* in the context of opioid-related behaviors. Using a multi-phase fentanyl IVSA paradigm, we found that *Reln* haploinsufficiency did not alter acquisition, extinction, or cue-induced reinstatement of fentanyl seeking. However, progressive ratio testing revealed a sex-dependent effect on motivational effort, with male *Reln*+/− mice exhibiting reduced motivation for fentanyl compared to male wild-type mice. In complementary non-contingent assays, the most prominent effect of reduced *Reln* expression emerged under conditions of acute pharmacological challenge where *Reln*+/− mice exhibited enhanced fentanyl-induced locomotion and elevated Fos immunoreactivity in the dorsal striatum.

### *Reln*-dependent modulation of acute fentanyl-induced locomotor and striatal activation

*Reln*+/− mice exhibited enhanced fentanyl-induced locomotion that was consistent across both doses tested and increased dorsal striatal Fos immunoreactivity following acute exposure at the lower dose of fentanyl (20 µg/kg). Importantly, no genotypic difference in *Fos* expression was detected in the dorsal striatum following cue-induced reinstatement of fentanyl seeking. These findings, together with the absence of changes during cue-induced reinstatement or CPP, suggests that *Reln* preferentially modulates acute pharmacological activation in the striatum rather than learned cue-driven neuronal activation.

These findings extend prior work linking Reelin signaling to drug-induced neuronal activation. Notably, similar phenotypes have been reported in psychostimulant models. De Guglielmo et al. (2022) demonstrated that *Reln* haploinsufficient mice exhibit enhanced cocaine-induced hyperlocomotion accompanied by increased *Fos* expression in the dorsal striatum, without a significant alteration in cocaine CPP. This parallel across two pharmacologically distinct drug classes suggests that *Reln* has a specific role in modulating drug-induced responses, separate from the learned associations that drive cue-induced drug seeking

### *Reln* haploinsufficiency does not alter opioid reinforcement

During fentanyl IVSA, *Reln*+/− mice exhibited unaltered acquisition, maintenance, and extinction of fentanyl self-administration, and total fentanyl intake did not differ by genotype during dose-response testing. Although nominally significant lever × genotype interactions emerged during dose-response testing, these did not survive post hoc correction, warranting caution in their interpretation. Similarly, during cue-induced reinstatement, while a significant session condition × genotype interaction was detected, genotypes did not differ significantly at either session condition after post hoc analysis. Fentanyl CPP was similarly unaffected. Together, these results indicate that *Reln* haploinsufficiency does not robustly alter the reinforcing effects of fentanyl or the expression of learned drug-seeking behavior. In contrast, the analysis of PR breakpoint revealed a sex-dependent role for Reelin signaling in the motivational effort required to obtain opioid reinforcement under conditions of increasing response cost, though this finding should be interpreted cautiously given the sample sizes within each subgroup.

Existing literature supports that *Reln*-dependent effects on drug-induced behavior are not uniform across drug classes, dose, and experimental approaches. Methamphetamine-induced hyperlocomotion was attenuated, rather than enhanced, in *Reln* haploinsufficient mice in a dose-dependent manner (Matsuzaki et al., 2007), a direction opposite to the fentanyl-induced locomotion observed here. Nucleus accumbens-specific *Reln* knockdown reduced cocaine CPP without altering cocaine-induced locomotion (Brida et al., 2025), a dissociation that contrasts with the present findings of enhanced fentanyl locomotion in the absence of altered CPP. This divergence may also reflect differences in developmental stages and regional specificity of *Reln* manipulation. The present study used germline *Reln* haploinsufficient mice, in which reduced *Reln* expression occurs throughout development and across all brain regions. In contrast, prior cocaine IVSA experiments used adult, region-specific knockdown confined to the nucleus accumbens of rats (Brida et al., 2025), making direct comparisons difficult.

Collectively, these findings indicate that the influence of Reelin signaling on drug-evoked behavior depends on the pharmacological mechanism of the drug, the brain region and cell type in which Reln is manipulated, and the developmental timing of the manipulation.

### Study Limitations

Several limitations of the present study warrant consideration. First, the use of germline *Reln* haploinsufficient mice precludes dissociation of developmental versus adult roles of Reelin signaling. Given the critical role of Reelin in neuronal migration, dendritic maturation, and synaptogenesis during development (D’Arcangelo et al., 1995; Ikeda & Terashima, 1997; Rice & Curran, 2001), it is possible that the locomotor and striatal-activation phenotypes reported here reflect developmental adaptations rather than an acute requirement for Reelin signaling in the adult brain. This interpretation is further complicated by the system-wide nature of the haploinsufficiency where reduced *Reln* expression occurs across all brain regions and cell types throughout the lifespan, making it impossible to attribute the observed effects specifically to dorsal striatal reelin signaling. Future studies using inducible time-, circuit- and cell-type-specific manipulations of *Reln* expression in adolescent or adult animals will be required to further isolate the contribution of Reelin signaling in the dorsal striatum to opioid-induced locomotion and neuronal activation.

Second, the Fos immunoreactivity results, while statistically significant, were obtained with a small cohort of male mice only and should be interpreted with appropriate caution. An independent replication in a larger, mixed-sex cohort will be necessary to confirm the robustness of this finding and to determine whether the enhanced Fos response generalizes across sexes.

Third, molecular analyses were restricted to a small panel of IEGs at a single timepoint, which do not capture all IEGs (Bahrami & Drabløs, 2016) nor the full transcriptional or epigenetic landscape of opioid exposure (Egervari et al., 2017; Gaddis et al., 2022). In particular, the absence of a genotypic difference in *Fos* expression during cue-induced reinstatement was assessed in a single brain region at a single timepoint; it remains possible that *Reln*-dependent differences in neuronal activation emerge in other brain regions, in other cell populations, or at different phases of the IVSA protocol.

Finally, although the present study detected significant genotype × sex interactions during both PR testing and cue-induced reinstatement, the study was not specifically designed or powered to fully characterize sex as a biological variable. The subgroup sample sizes within each sex-by-genotype combination limit the reliability and interpretability of these sex-stratified effects. Future studies specifically designed and powered to test sex as a biological variable will be necessary to determine whether *Reln*-dependent modulation of opioid reinforcement differs between males and females.

Collectively, these findings establish that Reelin preferentially modulates the acute pharmacological effects of opioids within striatal circuitry rather than the directing the associative learning processes that drive opioid reinforcement and drug-seeking. This dissociation positions *Reln+/− mice* as a valuable model for studying how Reelin signaling shapes neuronal responses to opioids, with implications for understanding the molecular mechanisms underlying opioid sensitivity.

## Authors contributions

Conceptualization: C.L., F.T.; Investigation: C.L., A.M.L., S.D., T.H., L.L., A.C.; Formal analysis: C.L.; Visualization: C.L.; Writing – original draft: C.L.; Writing – review & editing: F.T., C.L., S.D.; Supervision: F.T.

## Acknowledgments

We thank members of the Telese laboratory for helpful discussions. We acknowledge the support of Havilah Taylor for technical assistance with mouse colony management. This work was supported by NIH R01DA056602 to F.T.; and NIGMS T32 Training Grant (2T32GM133351) and 1F31DA063338-01A1 to S.D.

## Figures and Legends

**Supplemental Figure 1. Sex-stratified behavioral and molecular responses across fentanyl IVSA phases in WT and *Reln*+/− mice. (A-B)** Active (solid line) and inactive (dashed line) lever presses (A) or infusions (B) across all IVSA sessions (Day 1-15), stratified by sex and genotype. (**C-D)** Total lever presses (**C**) or breakpoint (**D**) during the progressive ratio session, faceted by sex and genotype. (**E)** Active (solid line) and inactive (dashed line) lever presses during extinction phase (Extiction, Day 1-10), faceted by sex and genotype. (**F)** Comparison of active and inactive lever presses during the first 30 minutes of the last extinction session (Ext., Day 10) and during the subsequent 30-minute cue-induced reinstatement session (Rnst), faceted by sex and genotype. (**G)** Relative expression (2^-ΔCt) of *Fos, Arc, Egr1*, and *Npas4* expression in the dorsal striatum following cue-induced reinstatement, normalized to Gapdh; faceted by sex and genotype. Data are mean ± SEM. Female: WT n=5, *Reln*+/− n=8; Male: WT n=8, *Reln*+/− n=5.

**Supplemental Figure 2. Total fentanyl intake follows an inverted U-shaped dose-response curve in both WT and *Reln*+/− mice**. (**A-C)** Total fentanyl intake (ug) averaged across the two sessions at each unit dose during the dose response phase for WT (n=13, black) and *Reln*+/− (n=13, blue) mice (**A**) or separated by females (**B;** WT n=5, Reln+/− n=8) and males (**C;** WT n=8, Reln+/− n=5). Data are mean ± SEM.

**Supplemental Figure 3. Sex-stratified fentanyl CPP and fentanyl-induced locomotion. (A)** Change in percent time spent in the fentanyl-paired room on the test day relative to baseline, for mice conditioned at 20 µg/kg or 80 µg/kg, faceted by sex and genotype. (**B)** Average locomotor speed (total distance / total time) during the first CPP conditioning session following acute fentanyl injection at 20 or 80 µg/kg, faceted by sex and genotype. Data are mean ± SEM. Female: WT n=4; *Reln*+/− n=4; Male: WT n=4, *Reln*+/− n=4.

