## Supplementary figures and images for "Reln haploinsufficiency enhances fentanyl-induced locomotion and striatal activity without affecting opioid reinforcement and relapse-like behavior"

### Supplementary Fig. 1

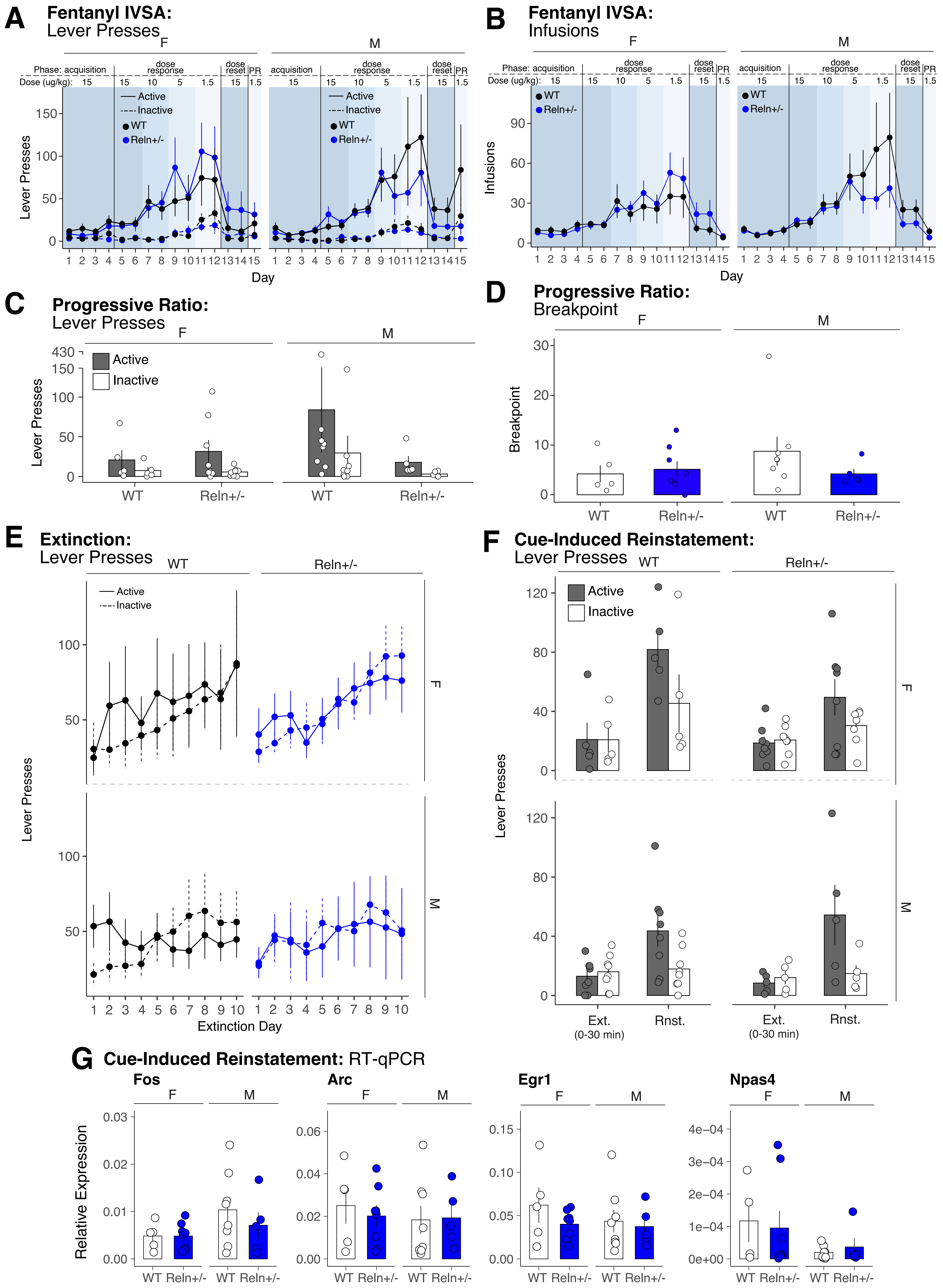

### Supplementary Fig. 2

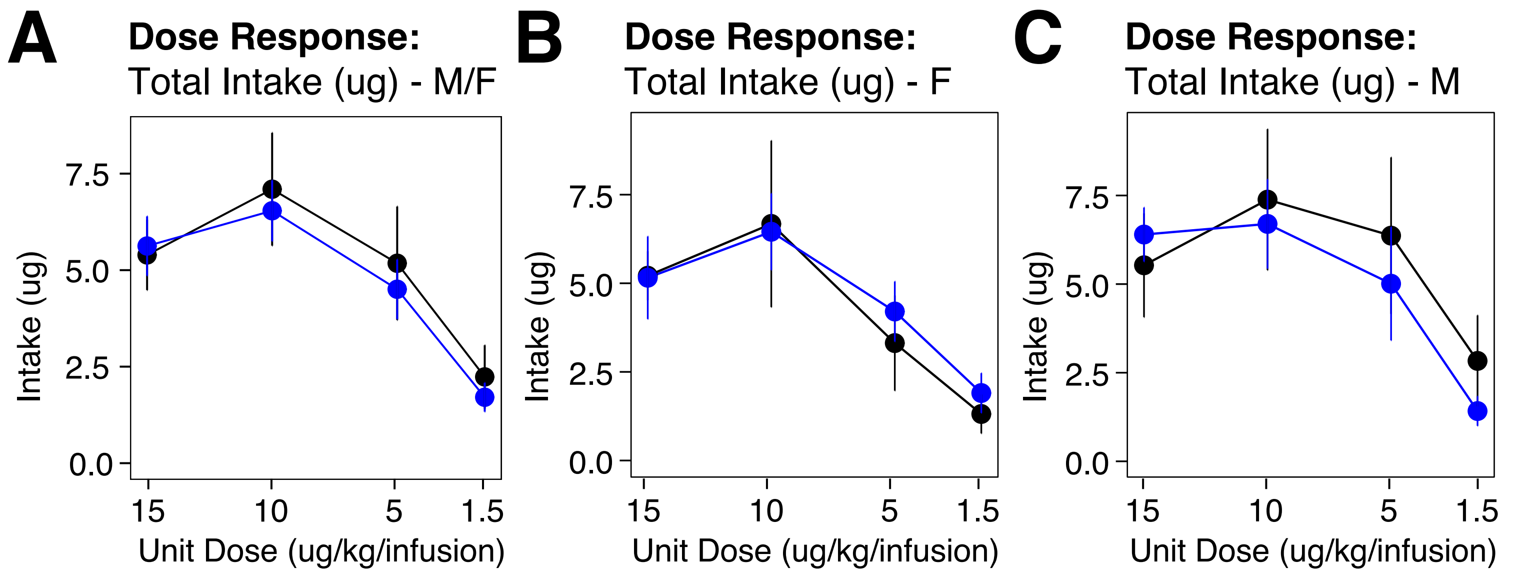

### Supplementary Fig. 3

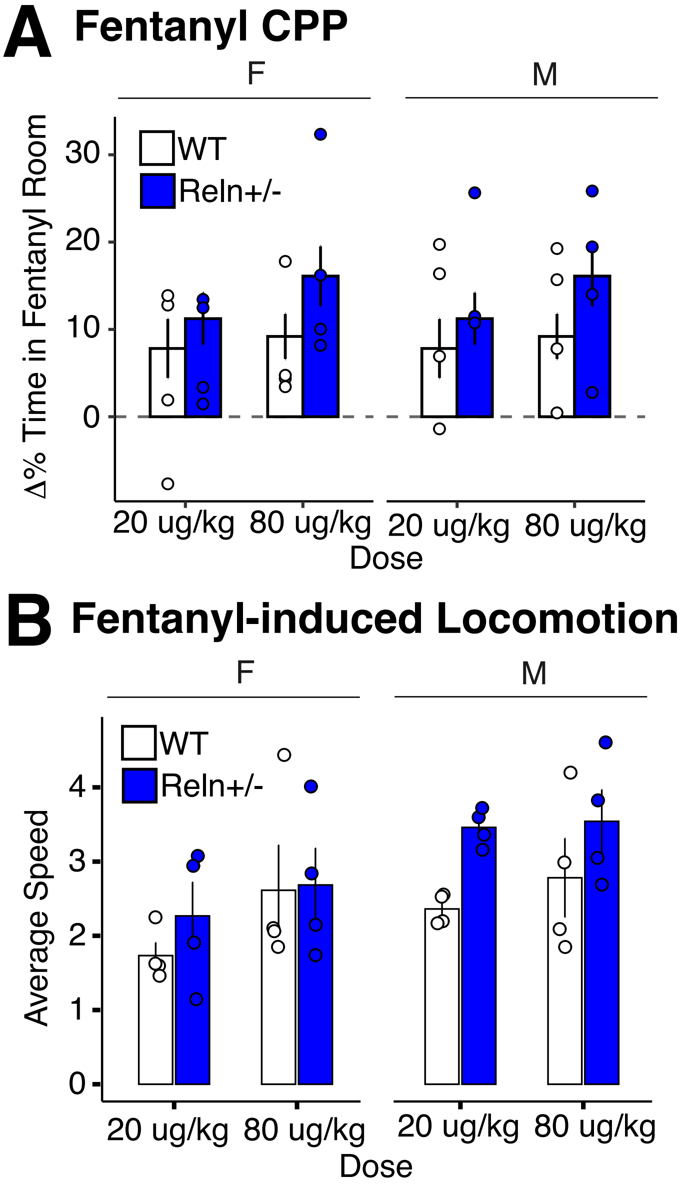
